# Values Encoded in Orbitofrontal Cortex Are Causally Related to Economic Choices

**DOI:** 10.1101/2020.03.10.984021

**Authors:** Sébastien Ballesta, Weikang Shi, Katherine E. Conen, Camillo Padoa-Schioppa

## Abstract

It has long been hypothesized that economic choices rely on the assignment and comparison of subjective values. Indeed, when agents make decisions, neurons in orbitofrontal cortex encode the values of offered and chosen goods. Moreover, neuronal activity in this area suggests the formation of a decision. However, it is unclear whether these neural processes are causally related to choices. More generally, the evidence linking economic choices to value signals in the brain remains correlational. We address this fundamental issue using electrical stimulation in rhesus monkeys. We show that suitable currents bias choices by increasing the value of individual offers. Furthermore, high-current stimulation disrupts both the computation and the comparison of subjective values. These results demonstrate that values encoded in orbitofrontal cortex are causal to economic choices.

## Introduction

In the 18^th^ century, Daniel Bernoulli, Adam Smith and Jeremy Bentham proposed that economic choices rely on the computation and comparison of subjective values (Niehans, 1990). This hypothesis continues to inform modern economic theory (Kreps, 1990) and research in behavioral economics (Kahneman and Tversky, 2000), but behavioral measures are ultimately not sufficient to prove the proposal (Camerer et al., 2005). Consistent with the hypothesis, when agents make choices, neurons in the orbitofrontal cortex (OFC) encode the subjective value of offered and chosen goods (Padoa-Schioppa and Assad, 2006). Moreover, neuronal activity in this area suggests the formation of a decision (Rich and Wallis, 2016). However, it is unclear whether these neural processes are causally related to choices. More generally, the evidence linking economic choices to value signals in the brain (Roesch and Olson, 2007; Bartra et al., 2013; Schultz, 2015) remains correlational(Stalnaker et al., 2015). Here we show that neuronal activity in OFC is causal to economic choices. We conducted two experiments using electrical stimulation in rhesus monkeys. Low-current stimulation increased the subjective value of individual offers and thus predictably biased choices. Conversely, high-current stimulation disrupted both the computation and the comparison of subjective values and thus increased choice variability. These results demonstrate a causal chain linking subjective values encoded in the OFC to valuation and choice. They set the stage to understand the mechanisms of economic decisions and goal-directed behavior, and to shed light on disorders affecting choices, including depression and drug addiction (Volkow and Li, 2004; Heyman, 2009).

Several lines of evidence link economic choices to the OFC. When subjects make choices, different groups of neurons encode the identities and values of offered and chosen goods (Padoa-Schioppa and Assad, 2006). Value encoding cells integrate multiple dimensions that characterize the options available in any given context (Roesch and Olson, 2005; Hare et al., 2008; Kennerley et al., 2009; Pastor-Bernier et al., 2019). Moreover, trial-by-trial variability in the activity of each group of neurons correlates with variability in choices (Padoa-Schioppa, 2013; Conen and Padoa-Schioppa, 2015). Computational models show that the cell groups identified in OFC are sufficient to generate binary choices (Rustichini and Padoa-Schioppa, 2015; Friedrich and Lengyel, 2016; Song et al., 2017; Zhang et al., 2018), and the population dynamics is consistent with decision making (Rich and Wallis, 2016). These results suggest that economic decisions might be formed within the OFC (Padoa-Schioppa and Conen, 2017), but causality has not been established. In principle, causal links between a neuronal population and a decision process are demonstrated if one can predictably bias choices using electrical stimulation (Cohen and Newsome, 2004; Clark et al., 2011). Building on this concept, classic work established the causal role of the middle temporal (MT) area in motion perception by showing that low-current stimulation biases (Salzman et al., 1990) while high-current stimulation disrupts (Murasugi et al., 1993) perceptual decisions. One challenge in using this approach for economic decisions is the lack of columnar organization in the OFC. Since neurons associated with different goods available for choice are physically intermixed (Conen and Padoa-Schioppa, 2015), it is impossible to selectively activate neurons associated with one particular good using electrical stimulation. Here we developed two experimental paradigms to circumvent this challenge.

## Results

In Exp.1, we examined whether perturbing OFC would disrupt choices. Monkeys chose between two juices labeled A and B (with A preferred) offered in variable amounts. The two offers were presented sequentially in the center of a computer monitor (**Fig.1A**). Trials in which juice A was offered first and trials in which juice B was offered first were referred to as “AB trials” and “BA trials”, respectively. An “offer type” was defined by two juice quantities in given order (e.g., [1A:3B] or [3B:1A]). The terms “offer1” and “offer2” referred to the first and second offer, independent of the juice type and amount. After a delay following offer2, two targets appeared on the two sides of the fixation point, and monkeys indicated their choice with a saccade. For each pair of juice quantities, the sequential order of the two offers varied pseudo-randomly. On roughly half of the trials, high-current stimulation (≥100 μA) was delivered in OFC for 0.5 s during offer1 presentation or during offer2 presentation (in separate sessions). In each session, trials with and without stimulation were pseudo-randomly interleaved (see **Methods**).

**Figure 1.**
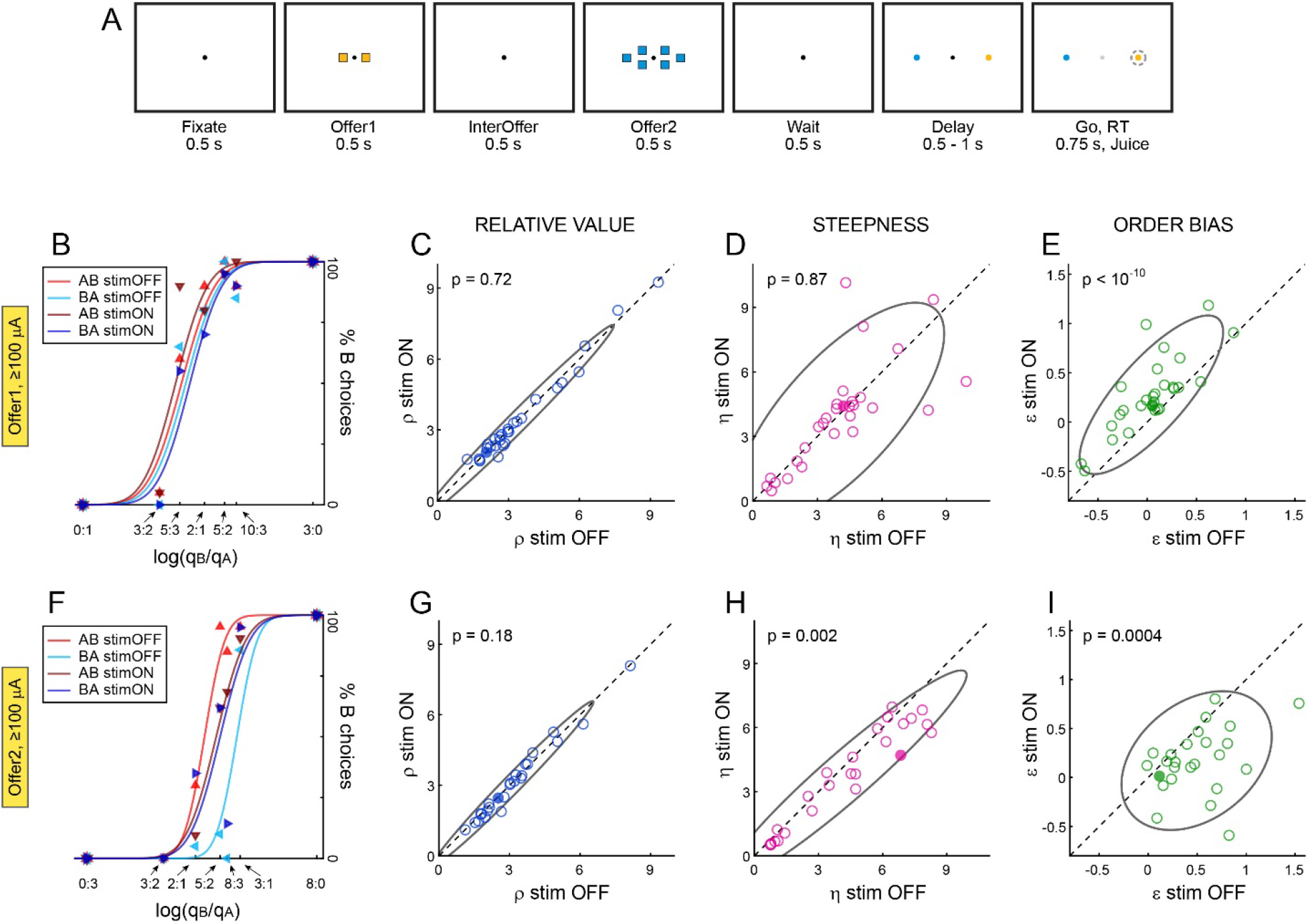
High-current perturbation of OFC disrupts valuation and decision. **A.** Experiment 1, design. Offers were represented by sets of squares, each color was associated with a juice type, and the number of squares represented the juice amount. In this trial, the animal chose between 2 drops of grape juice and 6 drops of peppermint tea. Offers were presented centrally and sequentially. After a delay, two targets associated with the two juices appeared on the two sides of the fixation point, and the animal indicated its choice with a saccade. Electrical stimulation was delivered in OFC during offer1 or offer2 (in separate sessions). **B.** Example session 1. In half of the trials, we delivered 125 μA current during offer1. The panel illustrates the choice pattern for AB trials (red) and BA trials (blue), separately for stimOFF trials (light) and stimON trials (dark). Data points are behavioral measures and lines are from probit regressions (Eq.1). In each condition (stimOFF, stimON), the order bias (ε) quantified the distance between the two flex points. In stimOFF trials, a small order bias favored offer2 (ε_stimOFF_ = 0.02). In stimON trials, the order bias increased (ε_stimON_ = 0.07). Hence, stimulation biased choices in favor of offer2. **CDE.** Population results for stimulation during offer1 (N = 29 sessions, ≥100 μA). Stimulation did not affect relative values (C); it did not consistently affect the sigmoid steepness (D); and it biased choices in favor of offer2 (E). **F.** Example session 2. Here 125 μA current was delivered during offer2. Stimulation induced a bias in favor of offer1 (ε_stimON_<ε_stimOFF_) and increased choice variability (shallower sigmoids in stimON trials; η_stimON_<η_stimOFF_). **GHI.** Population results for stimulation during offer2 (N = 25 sessions, ≥100 μA). Stimulation did not affect relative values (G); it reduced the sigmoid steepness (H); and it biased choices in favor of offer1 (I). In panels CDEGHI, filled symbols represent individual sessions in panels B and F and ellipses indicate 90% confidence intervals. All p values are from Wilcoxon tests, and very similar results were obtained using t tests.

For each group of trials (stimON, stimOFF), choice patterns were analyzed with a probit regression:

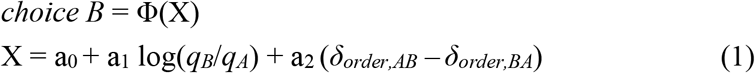

where *choice B* = 1 if the animal chose juice B and 0 otherwise, Φ was the cumulative distribution function of the standard normal distribution, *q_J_* was the quantity of juice J offered, and *δ*_*order,JK*_ = 1 in JK trials and 0 in KJ trials. From the fitted parameters, we derived measures for the relative value ρ = exp(−a_0_/a_1_), the sigmoid steepness η = a_1_, and the order bias ε = a_2_/a_1_. Intuitively, ρ was the quantity that made the animal indifferent between 1A and ρB, η was inversely related to choice variability, and ε was a bias favoring the first or second offer. Specifically, ε<0 (ε>0) indicated a bias in favor of offer1 (offer2).

In one representative session, electric current was delivered during offer1. The stimulation induced a choice bias in favor of offer2 (**Fig.1B**). This effect was consistent across N = 29 sessions: high-current stimulation during offer1 did not systematically alter the relative value or the sigmoid steepness, but it induced a systematic bias in favor of offer2 (**Fig.1CDE**). In a different set of N = 25 sessions, we delivered ≥100 μA during offer2. In this case, stimulation induced a systematic bias in favor of offer1 (**Fig.1FI**). These complementary effects are interpreted as high-current stimulation disrupting or interfering with the ongoing computation of the offer value, resulting in a choice bias for the other offer. In addition, stimulation during offer2, but not during offer1, significantly increased choice variability (**Fig.1FH**). This effect is interpreted as high-current stimulation disrupting value comparison (i.e., the decision), which took place upon presentation of offer2.

We also examined the effects of stimulation at lower currents. In essence, the effects observed at ≥100 μA diminished and gradually vanished when electric current was reduced to 50 μA and 25 μA (**Fig.2**). In summary, high-current stimulation of OFC disrupted the neural processes underlying economic choice, namely value computation during offer1, and value computation and value comparison during offer2.

**Figure 2.**
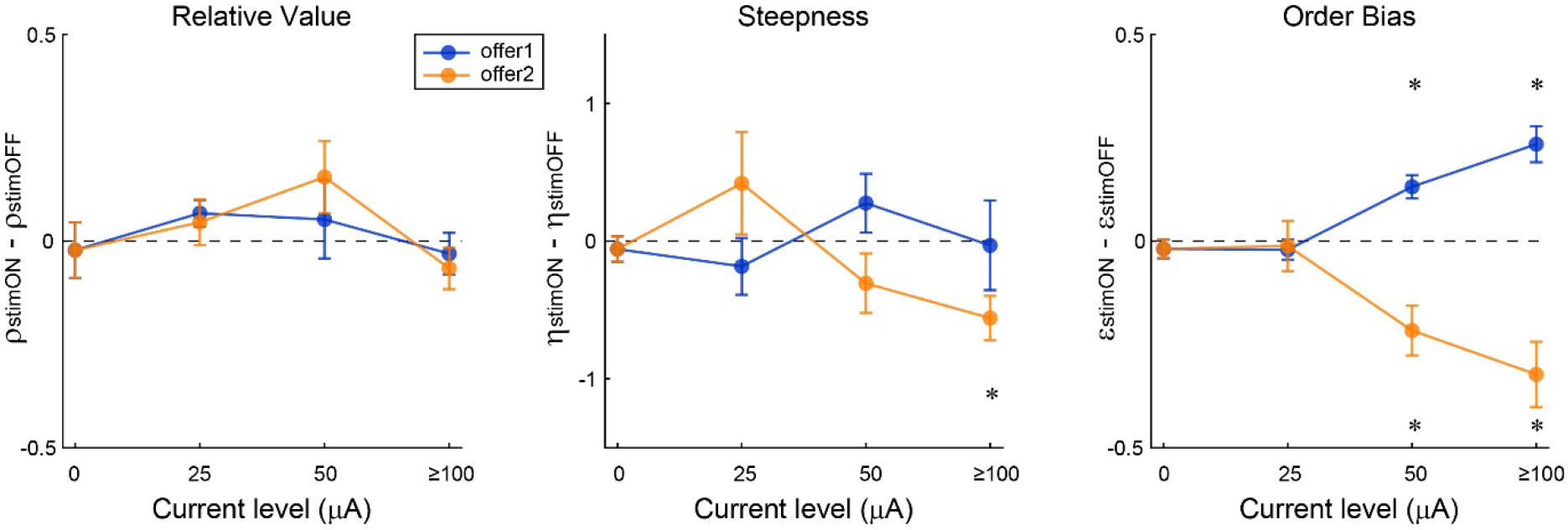
Effects of electrical stimulation at different current levels. We repeated Exp.1 varying current levels across sessions. Data shown here include N = 29/23/29 sessions in which 25/50/≥100 μA were delivered during offer1, N = 17/21/25 sessions in which 25/50/≥100 μA were delivered during offer2, and N = 50 control sessions (0 μA; 194 sessions total). The three panels illustrate the results obtained across sessions for the relative value, the sigmoid steepness and the order bias. In each panel, blue and yellow refer to stimulation during offer1 and offer2, respectively. Data points indicate averages across sessions and error bars indicate SEM. Asterisks highlight measures that differed significantly from zero (all p<0.003, Wilcoxon test). All other measures were statistically indistinguishable from zero (all p>0.05, Wilcoxon test). Statistical analyses based on t tests provided very similar results.

The results described so far showed that OFC perturbation disrupts valuation and choice. We next examined whether subjective values may be increased through physiological facilitation (Wolff and Olveczky, 2018). In Exp.2, we took advantage of the fact that neurons in OFC undergo range adaptation (Padoa-Schioppa, 2009; Kobayashi et al., 2010). **Fig.3** illustrates our rationale. In this experiment, monkeys chose between two juices offered simultaneously (**Fig.3A**). In these conditions, two groups of cells in OFC encode the offer values of juices A and B (Padoa-Schioppa and Assad, 2006; Padoa-Schioppa, 2013). Importantly, their tuning curves are quasi-linear and the gain is inversely proportional to the value range (range adaptation) (Padoa-Schioppa, 2009; Rustichini et al., 2017). Moreover, cells in each group adapt to their own value range. The effect of low-current stimulation is deemed equivalent to “artificially” increasing the firing rate of neurons in proximity of the electrode (Salzman et al., 1990; Histed et al., 2009). In turn, this increase in firing rate is equivalent to a small increase in the offer values. By virtue of range adaptation, for a given current, the increase in value is proportional to the value range (**Fig.3BC**). If an equal number of offer value A cells and offer value B cells are close to the electrode tip, then the effect of the electric current is equivalent to increasing both offer values. Crucially, if the two value ranges are unequal, the increases in offer values are also unequal. More specifically, the offer value of the juice with the larger range increases more. Hence, the net effect on choices is expected as follows: Low-current electrical stimulation should bias choices in favor of the juice offered with the larger value range (**Fig.3d; Supplementary Note**).

**Figure 3.**
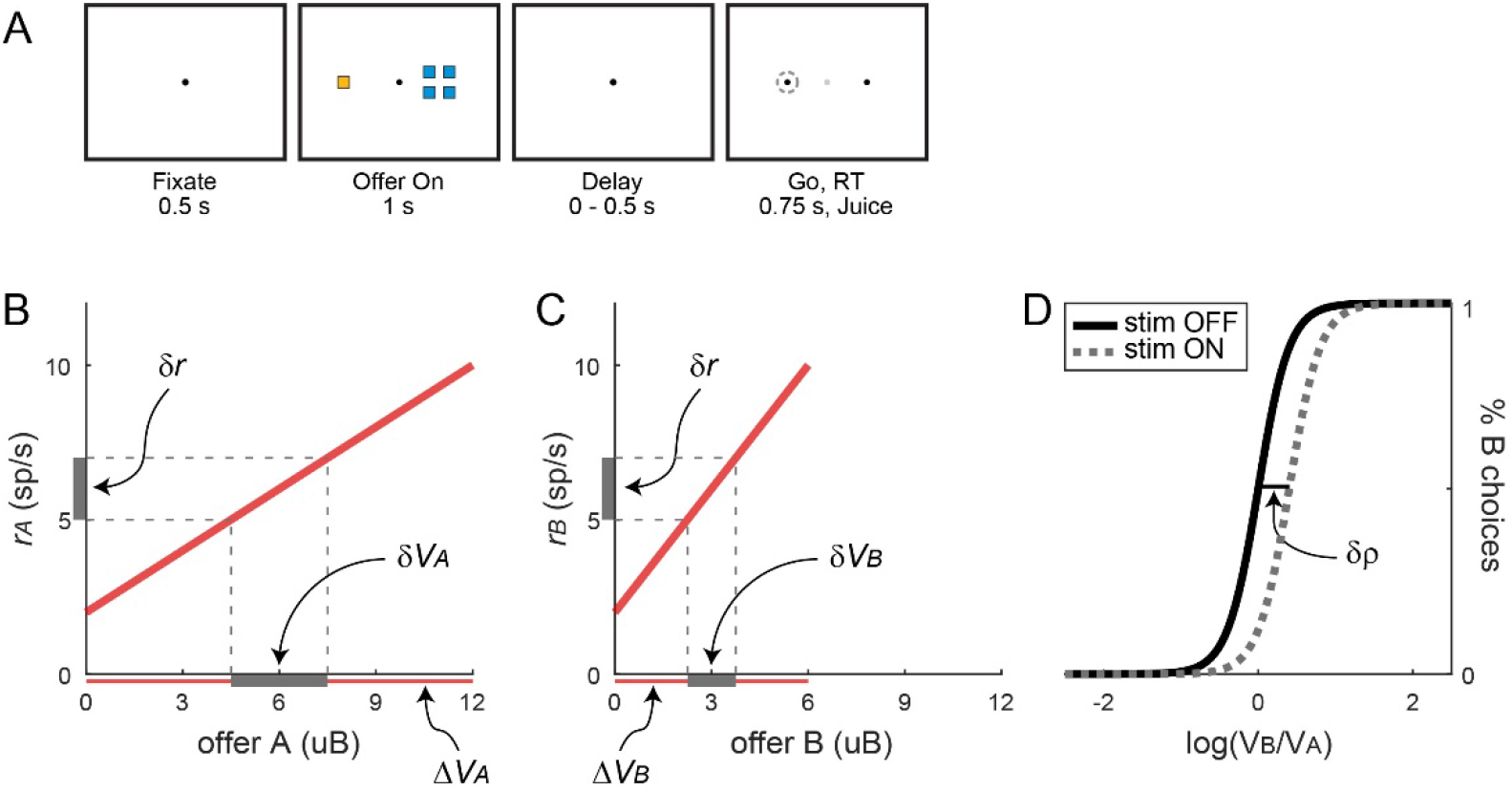
Prediction of range-dependent choice bias induced by electrical stimulation (facilitation). **A.** Experiment 2, design. Two offers are presented simultaneously. After a brief delay, the animal indicates its choice with a saccade. Electrical stimulation (50 μA) is delivered throughout offer presentation. **BCD.** Predictions for one example session. Panels B and C represent the (mean) tuning curves for pools of offer value A cells and offer value B cells under adapted conditions. Firing rates (y-axis) are plotted as a function of the offer values (x-axis) expressed in units of juice B (uB). Red horizontal lines represent the two value ranges, with ΔV_A_ > ΔV_B_. The same firing rate interval δr corresponds to different value intervals, with δV_A_ > δV_B_. Panel D represents choice patterns. Electrical stimulation increases both offer values, but the net effect is a choice bias in favor of juice A (δρ > 0). Conversely, in sessions where ΔV_A_ < ΔV_B_, δr induces δV_A_ < δV_B_, and electrical stimulation biases choices in favor of juice B (δρ < 0, not shown). See **Supplementary Note**.

We tested this prediction in two animals (see **Methods**). In each session, we selected juice types and quantity ranges such that value ranges (ΔV_A_, ΔV_B_) would differ. Electrical stimulation (50 μA) was delivered in OFC for 1 s during offer presentation. Trials with and without stimulation were pseudo-randomly interleaved. In each session, choice patterns were analyzed with a probit regression:

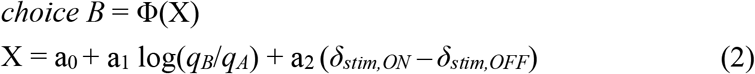

where *δ*_*stim,ON*_ = 1 in stimulation trials and 0 otherwise, and *δ*_*stim,OFF*_ = 1 − *δ*_*stim,ON*_. We computed the relative value for each group of trials and we defined the change in relative value induced by the stimulation as δρ = ρ_stimON_ − ρ_stimOFF_.

In one representative session, value ranges were such that ΔV_A_<ΔV_B_. Consistent with our prediction, electrical stimulation induced a bias in favor of juice B (δρ<0, **Fig.4A**). In another session, where ΔV_A_>ΔV_B_, we measured δρ>0 (**Fig.4B**). A population analysis found that the choice bias δρ and the difference in value range ΔV_A_−ΔV_B_ were strongly correlated across sessions. This result held true in each monkey (**Fig.4CD**). Data collected in paired sessions confirmed that range-dependent biases were not dictated by the juice types or the electrode position (**Fig.S1**).

**Figure 4.**
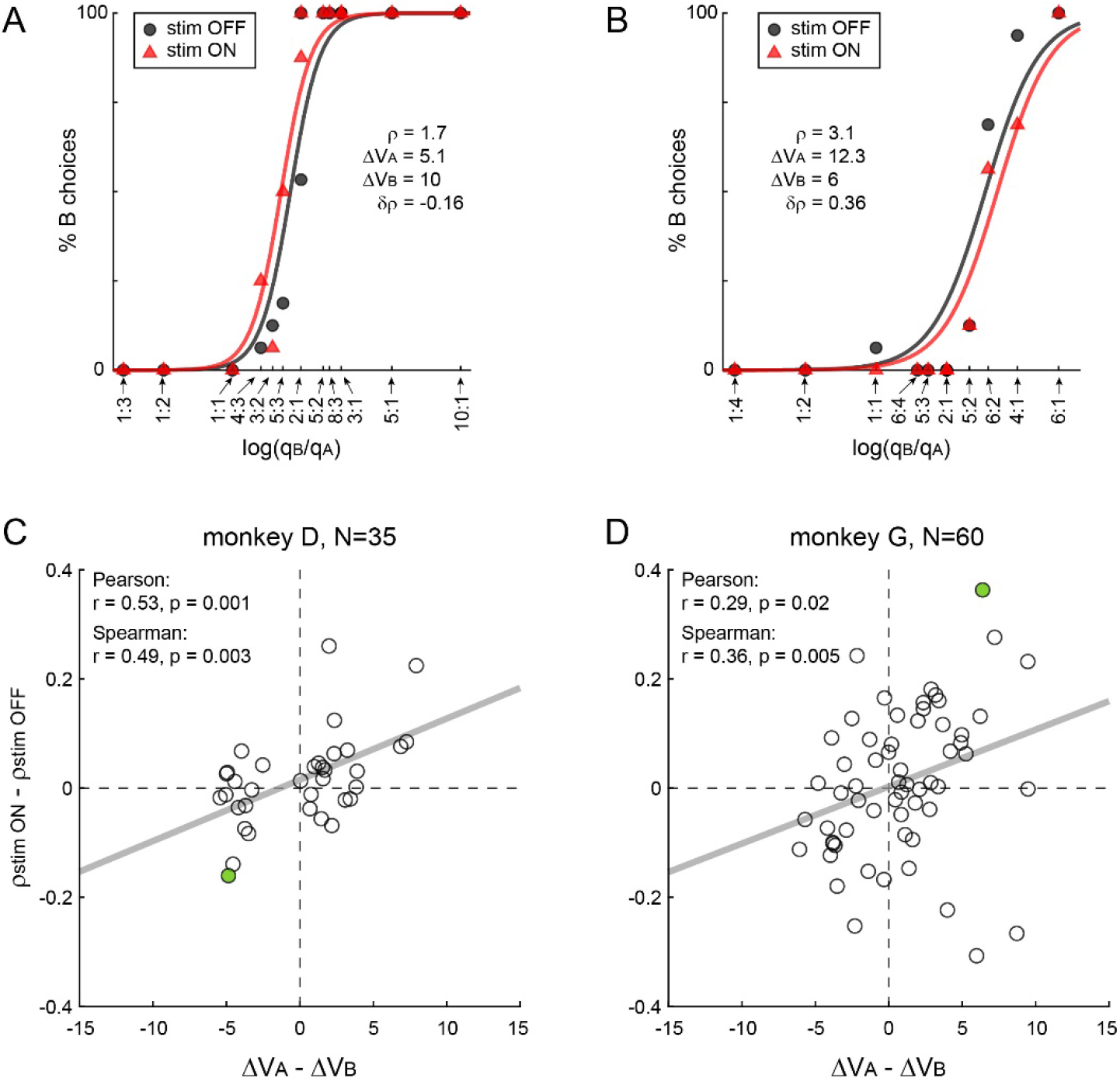
Range-dependent choice biases induced by neuronal facilitation of OFC. **A.** Example session 1. In this session, we set ΔV_A_ < ΔV_B_. Consistent with the prediction, electrical stimulation biased choices in favor of juice B (δρ < 0). **B.** Example session 2. In this case, we set ΔV_A_ > ΔV_B_. Electrical stimulation biased choices in favor of juice A (δρ > 0). **CD.** Population analysis. The two panels refer to the two animals. In each panel, the choice bias (δρ, y-axis) is plotted against the difference in value range (ΔV_A_−ΔV_B_, x-axis). Each data point represents one session, and the gray line is from a linear regression. Value ranges are expressed in units of juice B (uB). The two measures are significantly correlated in both monkey D (r = 0.53, p = 0.001, Pearson correlation test; r = 0.49, p = 0.003, Spearman correlation test) and monkey G (r = 0.29, p = 0.02, Pearson correlation test; r = 0.36, p = 0.005, Spearman correlation test). Data points highlighted in green are from the sessions illustrated in panels A and B.

The rationale of Exp.2 rests on the assumption that low-current stimulation increases the value of the two offers (**Fig.3**). An analysis of response times (RTs) supported this point. Under normal conditions (stimOFF), RTs decreased as a function of the chosen value. Electrical stimulation generally reduced RTs. Furthermore linear regressions of RTs onto the chosen value showed that this reduction was driven by lower offsets as opposed to steeper slopes (**Fig.S2**). These changes in RTs are as predicted if stimulation increases the subjective value of the chosen goods.

The results of Exp.2 were replicated in a secondary analysis of data from Exp.1. For this analysis, we pooled all trials (AB and BA) and all sessions (stimulation during offer1 or offer2), and we repeated the analysis conducted for Exp.2 (Eq.2). We found a significant correlation between the choice bias (δρ) and the difference in value range (ΔV_A_−ΔV_B_) when stimulation was delivered at 25 μA or 50 μA, but not when it was delivered at ≥100 μA (**Fig.S3**). The last observation indicated that the range-dependent facilitation observed at lower currents was fundamentally different from the disruption induced by high current. With respect to the effects of 50 μA stimulation, two aspects are noteworthy. First, 50 μA stimulation induced both facilitation and disruption (**Fig.S3B**; **Fig.2**, order bias). This finding is consistent with previous observations on area MT during perceptual decisions (Murasugi et al., 1993). Second, 50 μA stimulation induced range-dependent biases (**Fig.S3B**) but did not alter relative values on average (**Fig.2**, relative value). This finding is explained as follows. Electrical stimulation increased the value of both options, but more so for the option offered with a wider range. Thus the relative value changed as a function of the difference in value ranges. However, once sessions with ΔV_A_>ΔV_B_ and sessions with ΔV_A_<ΔV_B_ were pooled, there was no systematic change in relative value.

In the conditions examined here, different groups of neurons in OFC represent individual offer values, the binary choice outcome and the chosen value (Padoa-Schioppa, 2013; Ballesta and Padoa-Schioppa, 2019). Importantly, neurons encoding the binary choice outcome do not adapt to the value range, while chosen value cells adapt to the maximum range independent of the juice type (Conen and Padoa-Schioppa, 2019). Hence, physiological facilitation of these two cell groups is not expected to induce a choice bias for either juice depending on the value ranges. Thus range-dependent biases induced by stimulation are mediated by offer value cells (**Fig.3**).

## Discussion

In summary, it has long been hypothesized that economic choices rely on the assignment and comparison of subjective values (Niehans, 1990), but this proposal cannot be validated based on behavioral measures alone (Camerer et al., 2005). Consistent with the hypothesis, multiple brain regions represent values (Roesch and Olson, 2007; Schultz, 2015). However, value signals may inform a variety of mental functions including perceptual attention, learning and emotion (Schultz, 2015). Hence, to demonstrate the hypothesis, it is necessary to show a causal chain linking some neural representation of value to overt choices. Previous work indicated that OFC inactivation disrupts choices (Camille et al., 2011; Kuwabara et al., 2020), but other studies found different results (Gardner et al., 2017). Most importantly, previous work did not examine whether economic values can be increased through physiological facilitation – a necessary criterion to establish causality (Wolff and Olveczky, 2018). Here we have shown that neuronal processes taking place in OFC are causal to economic choices. In particular, choice biases induced by high current and range-dependent biases induced by lower current demonstrate a causal chain linking economic choices to subjective values encoded in OFC. In addition, the increase of choice variability observed in Exp.1 (upon stimulation during offer2 but not during offer1) indicates that OFC is directly involved in value comparison (i.e., the decision).

Electrical stimulation has often been used to assess causal relationships between neural substrates and behaviors (Cohen and Newsome, 2004; Clark et al., 2011). The experimental approach introduced here, combining neuronal adaptation and stimulation, may be used in future studies to assess causality for other cognitive functions controlled by the frontal lobe.

## Acknowledgments

This research was supported by the National Institutes of Health (grants number R01-DA032758 and R01-MH104494 to CPS and grant number F31-MH107111 to KEC) and by the McDonnell Center for Systems Neuroscience (pre-doctoral fellowship to WS). Author contributions: SB and WS collected and analyzed the data; KEC designed Exp.2; CPS supervised the project and wrote the manuscript. The authors have no competing interests. We thank H. Schoknecht for help with animal training and J. Assad, E. Bromberg-Martin, E. Fehr, D. Freedman, I. Monosov and L. Snyder for comments on the manuscript.

## Methods

All experimental procedures conformed to the NIH *Guide for the Care and Use of Laboratory Animals* and were approved by the Institutional Animal Care and Use Committee (IACUC) at Washington University.

The study was conducted on three male rhesus monkeys: G (age 8, 9.1 kg), J (age 7, 10.0 kg), and D (age 8, 11.5 kg). Before training, we implanted in each monkey a head-restraining device and an oval recording chamber under general anesthesia. The chamber (main axes, 50×30 mm) was centered on stereotaxic coordinates (AP 30, ML 0), with the longer axis parallel to a coronal plane. During the experiments, the animals sat in an electrically insulated enclosure with their head restrained. A computer monitor was placed 57 cm in front the animal. Behavioral tasks were controlled through custom-written software (http://www.monkeylogic.net/). The gaze direction was monitored by an infrared video camera (Eyelink; SR Research) at 1 kHz.

### Choice tasks

In Experiment 1 (Exp.1; monkeys G and J), animals chose between two juices labeled A and B, with A preferred. The two juices were offered sequentially and in variable amounts (**Fig1.A**). The terms “offer1” and “offer2” refer to the first and second offer, independently of the juice type and amount. Each trial began with the animal fixating a dot (0.35° of visual angle) in the center of the monitor. After 0.5 s, two offers appeared centrally and sequentially. Each offer was represented by a set of colored squares (side = 1° of visual angle), where the color indicated the juice type and the number of squares indicated the juice amount. Along with the offer, a small colored circle (0.75° of visual angle) appeared around the fixation dot. In the case of null offer (0 drops), the circle indicated to the animal the identity of the corresponding juice. The animal maintained center fixation throughout the initial fixation (0.5 s), offer1 time (0.5 s), inter-offer time (0.5 s), offer2 time (0.5 s), wait time (0.5 s), and delay time (0.5-1 s). At the end of the delay, the fixation point was extinguished and the animal indicated its choice with a saccade. It then maintained peripheral fixation for 0.6 s before juice delivery. Center fixation was imposed within 3°. Sessions typically included ~400 trials and offered quantities varied from trial to trial pseudo-randomly. We designed offer types such that in most trials the animal necessarily waited for offer2 before making a decision (Ballesta and Padoa-Schioppa, 2019). For each pair of juice quantities, the presentation order (AB, BA) and the spatial location of the saccade targets varied pseudo-randomly and were counterbalanced across trials. Stimulation was delivered in half of non-forced choice trials, pseudo-randomly selected.

In Experiment 2 (Exp.2; monkeys D and G), animals performed a similar task, except that the two juices were offered simultaneously (**Fig.3A**). After initial fixation (0.5 s), two offers appeared on the two sides of the fixation point. Offers remained on the monitor for 1 s and then disappeared. After a brief delay (0-0.5 s), the fixation point was extinguished and the animal indicated its choice with a saccade. The chosen juice was delivered after 0.75 s of peripheral fixation. Sessions typically included ~500 trials. Offered quantities and the spatial disposition varied from trial to trial pseudo-randomly. Previous work showed that in very similar conditions offer value cells in OFC undergo range adaptation (Padoa-Schioppa, 2009). Stimulation was delivered in roughly half of the trials, pseudo-randomly selected. We always tried to set the quantity ranges for the two juices such that the two value ranges would differ appreciably. However we could not fully control the difference in value ranges, because the relative value of the two juices (ρ) ultimately depended on the animal’s choices.

In some cases, we ran two paired sessions of Exp.2 back-to-back. In these cases, we left the stimulating electrode in place and we used the same two juices in both sessions, but we varied the quantity ranges such that the difference in value range ΔV_A_ − ΔV_B_ would be >0 in one session and <0 in the other session.

The quantity of juice associated with each square (quantum) was set equal to 70-100 μl in Exp.1, and to 75 μl in Exp.2 (the quantum always remained constant within a session). Across sessions, we used a variety of different juices associated with different colors, including lemon Kool-Aid (bright yellow), grape (bright green), cherry (red), peach (rose), fruit punch (magenta), apple (dark green), cranberry (pink), peppermint tea (bright blue), kiwi punch (dark blue), watermelon (lime) and 0.65 g/L salted water (light gray). This resulted in a large number of juice pairs.

### Electrical stimulation

The chamber provided bilateral access to OFC. Structural MRIs (1 mm sections) performed before and after surgery were used to guide electrode penetrations. Prior to the electrical stimulation experiments, we performed extensive neuronal recordings in each monkey using standard procedures (Ballesta and Padoa-Schioppa, 2019). Recordings and stimulation focused on the central orbital gyrus, in a region corresponding to area 13/11. The analysis of neuronal data confirmed that stimulation experiments focused on the same region examined in previous studies (Padoa-Schioppa and Assad, 2006; Ballesta and Padoa-Schioppa, 2019).

During stimulation sessions, low-impedance (100-500 kΩ) tungsten electrodes (100 μm shank diameter; FHC) were advanced using a custom-built motorized micro-drive (step size 2.5 μm) driven remotely. Stimulation trains were generated by a programmable analog output (Power 1401, Cambridge Electronic Design) and triggered through a TTL by the computer running the behavioral task. Electric currents were generated by an analog stimulus isolator (Model 2200, A-M Systems). The parameters used for electrical stimulation were as follows.

In Exp.1, electric current was delivered during offer1 or during offer2 (in separate sessions). Stimulation started 0-100 ms after offer onset and lasted 300-600 ms. The stimulation train was constituted of bipolar pulses (200 μs each pulse, 100 μs separation between pulses) delivered at 100-333 Hz frequency (Salzman et al., 1990; Murasugi et al., 1993; Merrill et al., 2005; Kim et al., 2015). In different sessions, current amplitudes varied between 25 and 125 μA. Stimulation was performed in both hemispheres of monkey G (left: AP 31:36, ML −7:−12; right: AP 31:36, ML 4:9) and in both hemispheres of monkey J (left: AP 31:35, ML −8:−10; right AP 31:35, ML 6:10). Our data set included a total 144 stimulation sessions and 50 control sessions (see **Table S1**). Electric current was delivered either unilaterally or bilaterally, in separate sessions. For each current level, the two groups of sessions were combined in the analysis. Analysis of the conditions for which we had two sizeable data sets (namely, offer1 stimulation) indicated that unilateral and bilateral stimulation had similar effects on choices.

In Exp.2, the stimulation train (bipolar pulses, 200 Hz frequency) was delivered throughout offer presentation, for 1 s. Stimulation was always unilateral, and current amplitude was always set at 50 μA. Stimulation was performed in the left hemisphere of monkey D (AP 31:36, ML −6:−10) and in the left hemisphere of monkey G (AP 31:36, ML −7:−11). Trials with stimulation (stimON) and without stimulation (stimOFF) were pseudo-randomly interleaved.

Electrical stimulation did not systematically alter error rates. In both experiments, errors were defined as fixation breaks occurring any time prior to trial completion. In Exp.1, error rates were not affected by stimulation in any of the experimental conditions (25 μA, offer1, p = 0.10; 25 μA, offer2, p = 0.68; 50 μA, offer1, p = 0.15; 50 μA, offer2, p = 0.88; ≥100 μA, offer1, p = 0.20; ≥100 μA, offer2, p = 0.46; Wilcoxon test, two animals combined). Similarly, stimulation did not alter error rates in Exp.2 (p = 0.87; Wilcoxon test, two animals combined).

### Data analysis

All analyses were conducted in Matlab (MathWorks Inc). For the primary analysis of data from Exp.1, choice patterns were analyzed with probit regressions, separately for stimOFF trials and stimON trials (Eq.1). For each group of trials, we derived measures for the relative value of the juices (ρ), the sigmoid steepness (η) and the order bias (ε). The effects of electrical stimulation on each parameter were assessed using Wilcoxon signed-rank tests and paired t tests (**Figs.1-2)**, **Fig.2**).

For data from Exp.2, we first ran two independent probit regressions for stimON trials and stimOFF trials. We found that electrical stimulation did not systematically alter the sigmoid steepness (**Fig.S4**). Thus we ran a probit regression assuming equal steepness for the two groups of trials (Eq.2). Except for **Fig.S4**, all the results presented here were obtained from the latter fit. Referring to Eq.2, we defined ρ_stimON_ = exp(−(a_0_+a_2_)/a_1_) and ρ_stimOFF_ = exp(−(a_0_−a_2_)/a_1_).

At the time of Exp.1, we had not planned to examine range-dependent choice biases. To examine these effects, we pooled sessions in which stimulation was delivered during offer1 or offer2, and we re-analyzed data using the same procedures used for Exp.2.

Because range-dependent choice biases were quantitatively small (**Fig.4**), it was important that the two measures of relative value obtained for each session be accurate and precise. Thus we imposed two criteria on the choice patterns: (a) saturation – for each juice, in at least one offer type, the animal should choose the same juice consistently (≥90% of trials); (b) incomplete separation – for at least two offer types, the animal should split decisions between the two options (<90% choices for each juice). The latter criterion ensured convergence of the probit fit. Only sessions that satisfied these criteria in both conditions (stimON, stimOFF) were included in the analysis. For Exp.2, we ran a total of 101 sessions (36 from monkey D, 65 from monkey G). The exclusion criteria removed 6 sessions (1 from monkey D, 5 from monkey G). Including these sessions in the analysis did not substantially alter the results. For Exp.1, the primary analysis included all sessions. For the analysis of range-dependent biases, the exclusion criteria removed 9 of 46 sessions at 25 μA, 9 of 44 sessions at 50 μA, and 20 of 54 sessions at ≥100 μA. Including these sessions in the analysis made the correlation between δρ and ΔV_A_−ΔV_B_ non-significant at 25 μA, but it did not substantially alter the results obtained at 50 μA and ≥100 μA (**Fig.S3**). Conversely, imposing these criteria on the primary analysis of Exp.1 data did not alter the results in any significant way.

## Supplementary Note: Range-dependent Effects of Electrical Stimulation

Here we formalize the prediction illustrated in **Fig.3**. As a premise, previous work found that the tuning curves of offer value cells in OFC are quasi-linear (Rustichini et al., 2017) and the proportion of neurons presenting positive versus negative encoding is roughly 3:1 (Padoa-Schioppa, 2013; Ballesta and Padoa-Schioppa, 2019). Importantly, cells in each group adapt to their own range, not to the maximum range (Conen and Padoa-Schioppa, 2019). Neurons associated with the two two juices (A and B) are physically interleaved (Conen and Padoa-Schioppa, 2015).

For given offers q_A_ and q_B_, r_A_ and r_B_ indicate the average firing rate for the two pools of offer value cells. The effect of stimulation (facilitation) is a small increase in this average firing rate, such that r_A_ → r_A_ + δr_A_ and r_B_ → r_B_ + δr_B_. Since the two neuronal populations are physically intermixed, electrical stimulation affects both of them equally. In other words, δr_A_ = δr_B_ = δr.

For each population, and for each juice type, a small increase in firing rate (δr) is equivalent to a small increase of offered value (δV_A_, δV_B_). Since offer value cells undergo range adaptation,

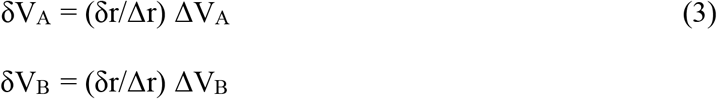

where Δr is the range of firing rates (which is the same for both juices), and ΔV_A_ and ΔV_B_ are the ranges of offered values (Padoa-Schioppa, 2009).

We aim to understand how electrical stimulation will affect choices – that is, how the relative value ρ will change under electrical stimulation. To do so, we write the conditions of choice indifference. We assume linear indifference curves and we indicate with V(J) = uJ the value of one unit (one quantum) of juice J. In the absence of stimulation:

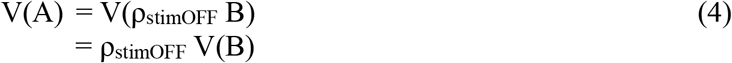

In the presence of stimulation:

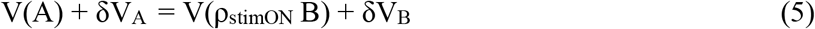

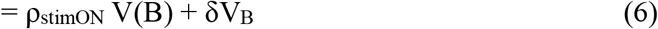

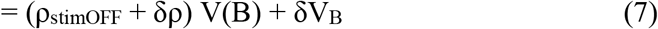

In the last passage, we defined δρ = ρ_stimON_ − ρ_stimOFF_. Now we substitute Eq.4 in Eq7 and we re-arrange:

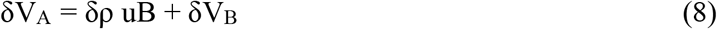

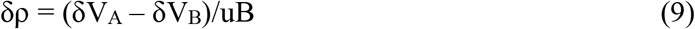

Finally, we substitute Eq.3 in Eq.9:

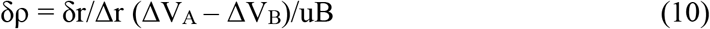

Eq.10 captures the key prediction: If decisions are primarily based on the activity of offer value cells, the net effect of electrical stimulation is to change the relative value of the juices by a quantity proportional to the difference in value ranges. Notably, by pooling sessions in **Fig.4** we effectively assumed that δr/Δr and uB remain constant across sessions. In practice, this might not be true because of variability in stimulation efficacy and because the subjective value of juice B might vary from session to session. These sources of variability effectively add noise to our measurements. However, the prediction that δρ and (ΔV_A_ − ΔV_B_) should have the same sign is not affected by these factors.

### Supplementary Table

**Table S1.** Data set for Exp.1. Labels uni/bi indicate unilateral/bilateral stimulation. For sessions indicated by ≥100 μA, current levels were set in the range 100-150 μA (typically at 125 μA).

| Stimulation interval | Current Level | Mode | Number of sessions |  |  |
| --- | --- | --- | --- | --- | --- |
|  |  |  | monkey G | monkey J | total |
| offer 1 | $\geq 100 \mu\text{A}$ | uni | 5 | 8 | 29 |
|  |  | bi | 15 | 1 |  |
| | 50 $\mu\text{A}$ | uni | 9 | 7 | 22 |
|  |  | bi | 4 | 3 |  |
| | 25 $\mu\text{A}$ | uni | 11 | 4 | 29 |
|  |  | bi | 14 | 0 |  |
| offer 2 | $\geq 100 \mu\text{A}$ | uni | 14 | 6 | 25 |
|  |  | bi | 3 | 2 |  |
| | 50 $\mu\text{A}$ | uni | 11 | 10 | 22 |
|  |  | bi | 0 | 0 |  |
| | 25 $\mu\text{A}$ | uni | 10 | 5 | 17 |
|  |  | bi | 2 | 0 |  |
| control | 0 $\mu\text{A}$ | -- | 30 | 20 | 50 |
| total | -- | -- | 131 | 63 | 194 |

**Figure S1.**
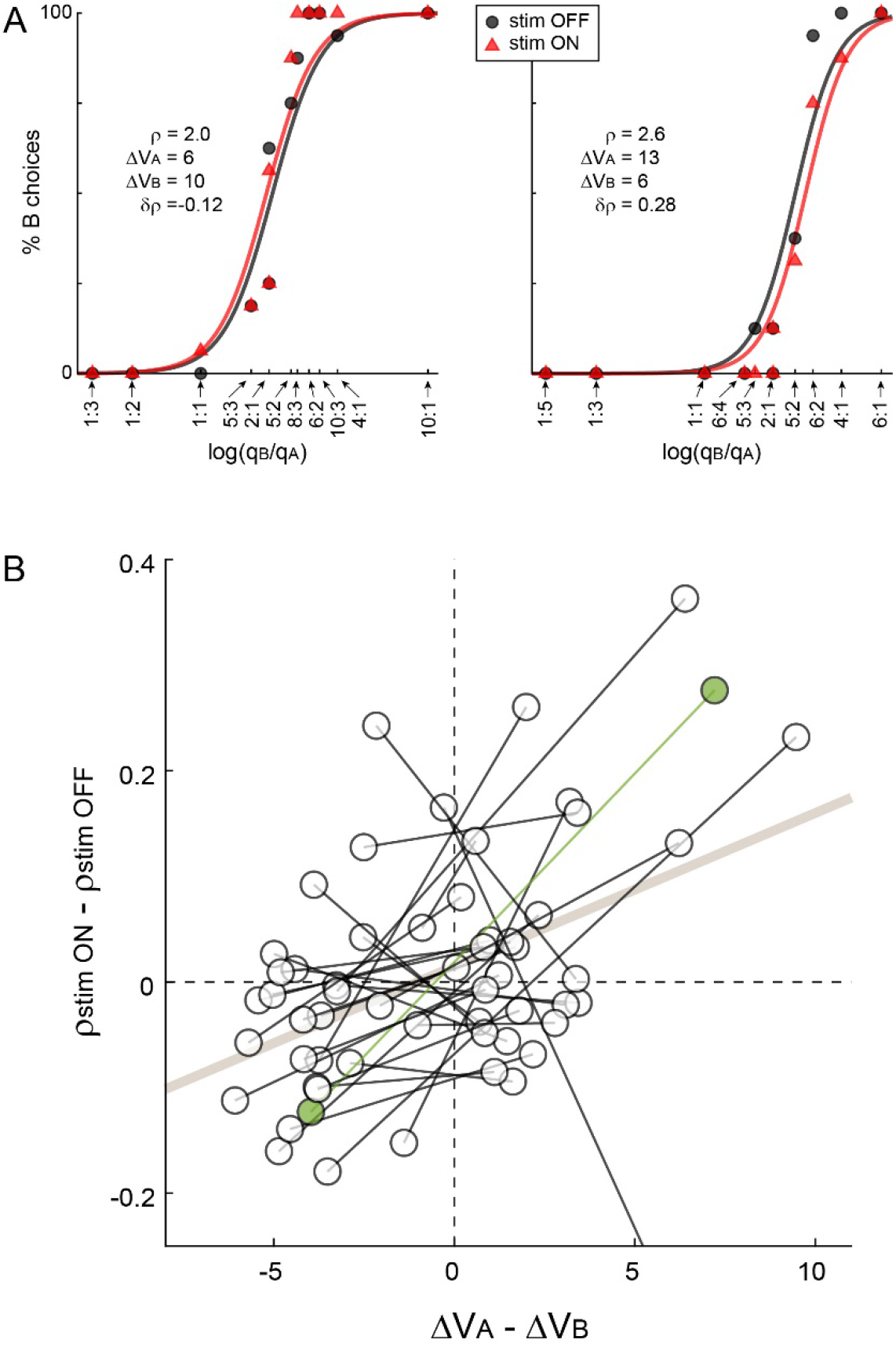
Exp.2, results obtained in paired sessions. In some case, we ran two back-to-back sessions offering the same two juices and leaving the electrode in place, but changing the quantity ranges such that ΔV_A_−ΔV_B_ would have opposite signs. **A.** Example of paired sessions. **B.** Population analysis (N = 29). Each pair of sessions in the scatter plot is connected by a line, of which we computed the slope. Data from the two monkeys are pooled. Across the population, slopes were typically >0 (p = 0.02, one-tailed Wilcoxon signed-rank test). Green data points correspond to the sessions in panel A. Hence, range-dependent biases were not dictated by the juice pair or by the location of the electrode within OFC.

**Figure S2.**
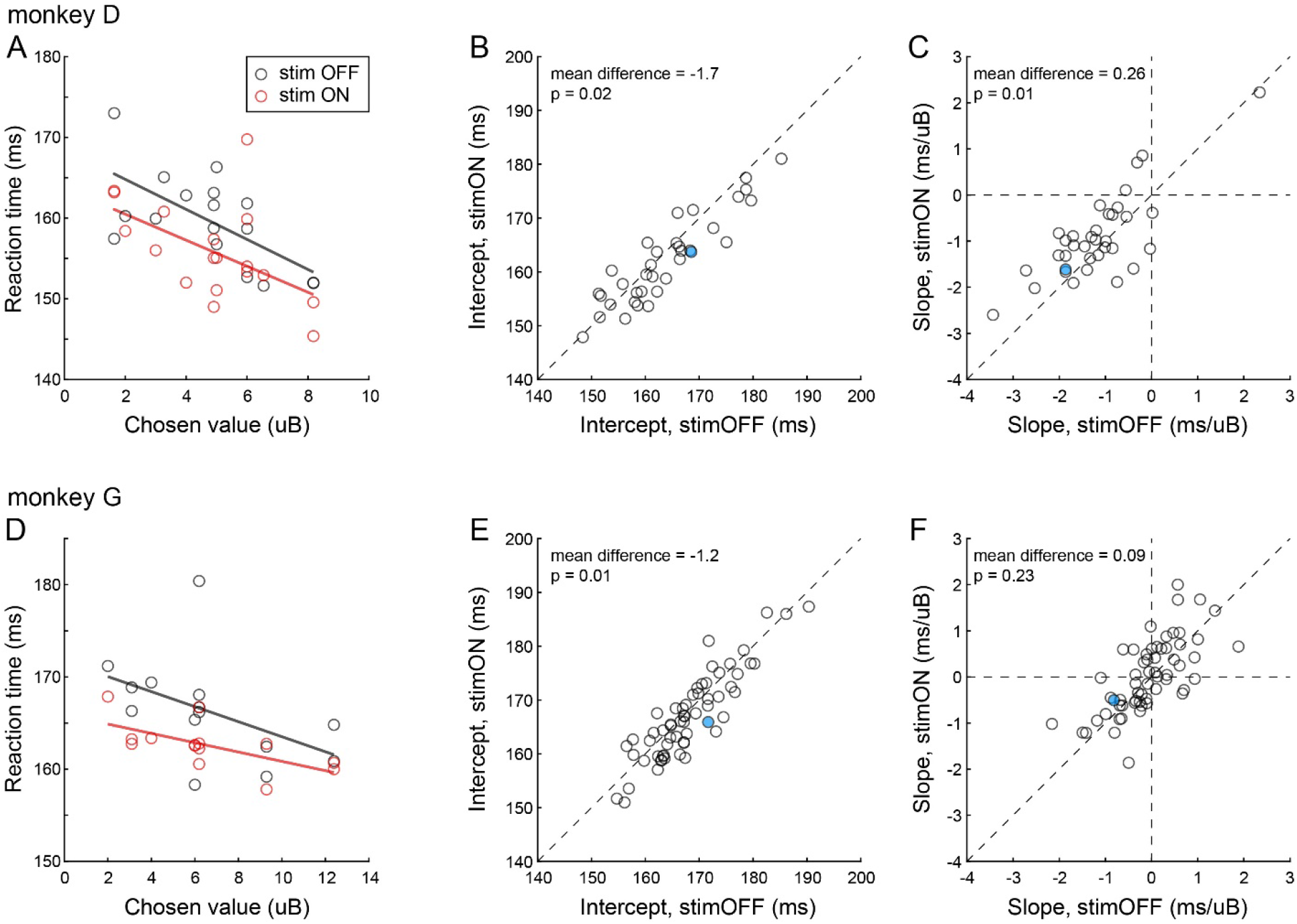
Exp.2, analysis of response times (RTs). **A.** Example session 1. Each data point represents one trial type and the two lines were obtained from linear regressions. Under normal conditions (stimOFF, black), RTs decreased as a function of the chosen value (x-axis). Electrical stimulation (stimON, red) generally reduced RTs. Linear fits reveal that lower RTs were due to a lower intercept, as opposed to a steeper (i.e., more negative) slope. **BC.** Population analysis, monkey D (N = 35). For each session, we regressed RTs onto the chosen value, separately for stimOFF and stimON trials. We then compared the intercepts and the slopes at the population level. The picture emerging from panel A was confirmed for the population. In panel B (intercept), each data point represents one session. The population is significantly displaced below the identity line (p<0.02, Wilcoxon test). In panel C (slope), it can be noticed that the slope under stimulation was shallower (less negative), probably due to a floor effect. Filled data points correspond to the session shown in panel A. **D.** Example session 2. Same format as in panel A. **EF.** Population analysis, monkey G (N = 60). Same format as in panels BC. Electrical stimulation significantly lowered the intercept but did not significantly alter the slope. Filled data points correspond to the session shown in panel D. In panels BCEF, values indicated in the insert refer to the difference between the stimON measure and the stimOFF measure, averaged across the population. All p values are from Wilcoxon tests, and t tests provided very similar results.

**Figure S3.**
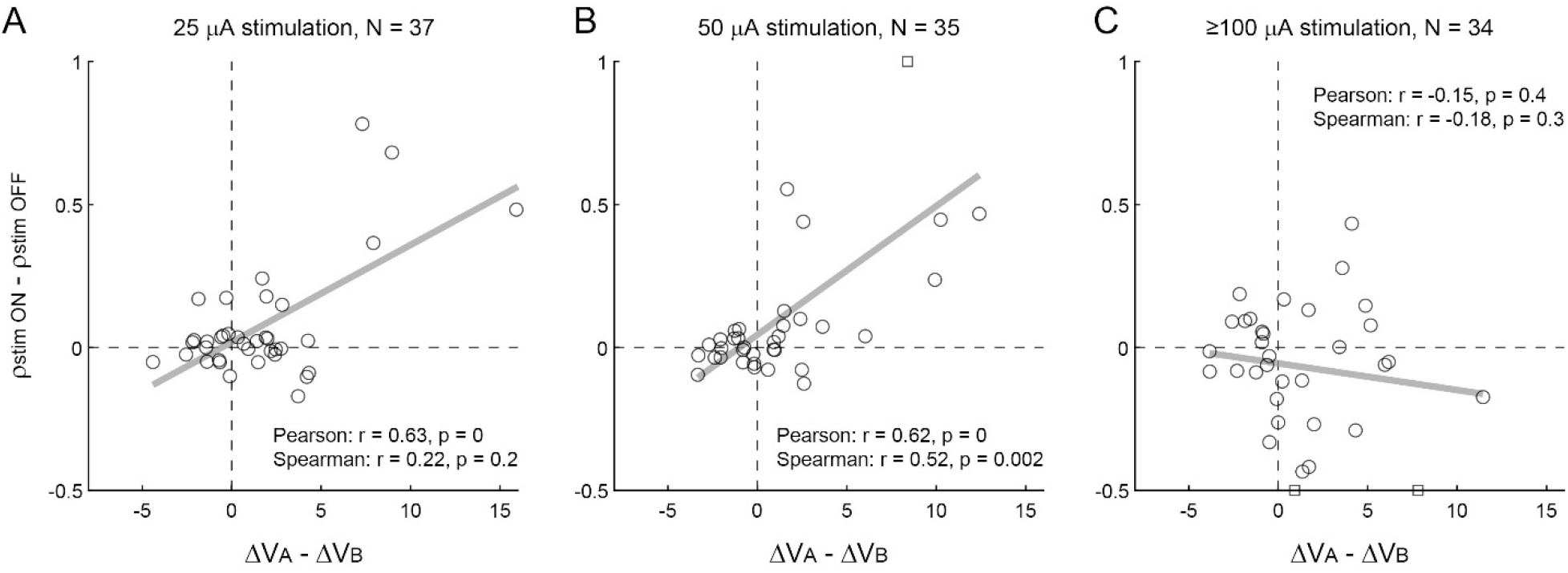
Exp.1, range-dependent choice biases. **ABC.** Results obtained when electric current was delivered at 25 μA, 50 μA and ≥100 μA. In each panel, x-and y-axes represent the difference in value range (in uB) and the difference in relative value, respectively. Each data point represents one session, and squares indicate outliers. Sessions from the two animals and with different stimulation times (offer1 or offer2) were pooled. Gray lines were obtained from linear regressions. Each panel indicates the p values obtained from Pearson and Spearman correlation tests. In essence, the choice bias imposed by the stimulation (δρ) was correlated with the difference in value ranges (ΔV_A_−ΔV_B_) at low current (25 μA; weakly) and intermediate current (50 μA), but not at high current (≥100 μA).

**Figure S4.**
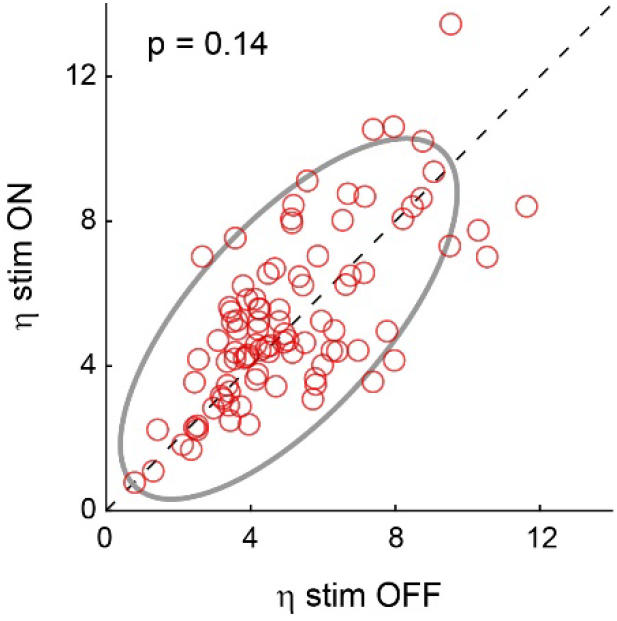
Stimulation in Exp.2 did not systematically alter the sigmoid steepness. For this analysis, the two groups of trials (stimOFF, stimON) were examined separately (see **Methods**). The two axes represent the sigmoid steepness in the two conditions. Sessions from the two animals were pooled, and each data point represents one session. The gray ellipse represents the 90% confidence interval. The p value is from a Wilcoxon test.

